# A novel mechanism of ceftolozane-tazobactam resistance in *Pseudomonas aeruginosa* mediated by L2 β-lactamase

**DOI:** 10.64898/2026.03.10.710737

**Authors:** Preeti Garai, Sophia H. Nozick, Caroline C. Jozefczyk, Hannah Nam, J. Nicholas O’Donnell, Egon A. Ozer, Alan R. Hauser, Nathaniel J. Rhodes

## Abstract

The prevalance of non-susceptibility to ceftolozane-tazobactam (C/T) among *Pseudomonas aeruginosa* remains low but novel mechanisms of C/T resistance are of concern. Herein, we describe a novel *Pseudomonas aeruginosa* genotype associated with high-level C/T resistance (>256/4 µg/mL) in a single patient. Whole genome sequencing of the isolate was compared to that of a susceptible isolate cultured from the same patient two months earlier. Analysis of the sequences revealed two different *P. aeruginosa* high-risk clones: ST111 followed by ST235. The C/T-resistant ST235 isolate contained five copies of a genetic element comprised of an L2 β-lactamase gene (*bla*_L2_) and a truncated *ampR*_L2_ transcriptional regulator gene, which are commonly found together in *Stenotrophomonas maltophilia* strains and have not been reported to mediate resistance to C/T. Comparative genomic analysis with other *P. aeruginosa* isolates failed to identify alternative explanations for the observed C/T resistance. We found that exogenous expression of *bla*_L2_ modestly increased C/T MICs in genetically distinct *P. aeruginosa* strains. A screen of our archived isolates identified two *P. aeruginosa* clinical isolates, PS2045 and PS2046, with one and two copies, respectively, of the genetic element containing *bla*_L2_ and truncated *ampR*_L2_. Interestingly, disruption of the gene *bla*_L2_ but not the truncated *ampR*_L2_ in PS2045 led to a significant decrease in C/T MIC. Thus, we report a novel mechanism of C/T resistance in *P. aeruginosa* mediated by an L2 β-lactamase independently of its canonical regulator AmpR_L2_.

## Introduction

Rising rates of antibiotic resistance in the bacterial pathogen *Pseudomonas aeruginosa* pose a serious threat to public health. Infections caused by antibiotic-resistant strains of *P. aeruginosa* are extremely difficult to treat and are associated with severe morbidity and high mortality rates. Ceftolozane-tazobactam (C/T), a combination of an antipseudomonal cephalosporin, ceftolozane, and a β-lactamase inhibitor, tazobactam (1, 2), has been efficacious in treating *P. aeruginosa* infections (2–5). The high activity of C/T against multi-drug resistant (MDR), extensive-drug resistant (XDR) and difficult to treat (DTR) strains of *P. aeruginosa* (2, 6–10) is attributable to the ability of C/T to evade several antibiotic resistance mechanisms employed by *P. aeruginosa* that inactivate first-line antipseudomonal antibiotics (11, 12).

The prevalence of *P. aeruginosa* non-susceptibility to C/T remains low among clinical isolates from critically ill patients, ranging from 3 to 18 percent (4, 13–15), but the evolution of C/T resistance during therapy is concerning. While the most common mechanism of C/T resistance in *P. aeruginosa* involves mutations leading to overexpression or increased hydrolytic activity of the class C β-lactamase AmpC (16–18), other mechanisms such as the acquisition of extended-spectrum β-lactamase (ESBL) genes and over-expression of efflux pumps have been occasionally reported (17, 19–24). Given the results of the recent multicenter CACTUS study of hospitalized patients in the USA, which found greater clinical efficacy (OR=2.07, 95% CI 1.16-3.70) with C/T vs. ceftazidime-avibactam (CZA) across a variety of *P. aeruginosa* infections (25), it is expected that C/T use and resistance will increase in the near term. Thus, it is crucial to understand emerging C/T resistance mechanisms in *P. aeruginosa*. Herein, we describe a novel mechanism of C/T resistance mediated by acquisition of the L2 β-lactamase gene (*bla*_L2_) by a *P. aeruginosa* strain cultured from a patient who had received multiple courses of C/T and other antimicrobials.

## Materials and Methods

### Patient case and clinical course

All clinical data were obtained from the Northwestern Memorial Hospital (NMH), Chicago electronic medical record. A case report exemption was obtained from the Northwestern University and Midwestern University Institutional Review Boards.

A 51-year-old male with a history of quadriplegia, autonomic hyperreflexia, neurogenic bladder and chronic foley catheterization complicated by *Enterococcus faecalis* urinary tract infections (UTIs), decubitus ulcers complicated by chronic osteomyelitis, chronic respiratory failure with tracheostomy complicated by recurrent *P. aeruginosa* pneumonia, and *Clostridioides difficile* infection presented from a skilled nursing facility with reports of low oxygen saturations, increased respiratory secretions, and hyponatremia. Six months prior to the current admission, the patient was admitted with multidrug-resistant *P. aeruginosa* pneumonia and was treated with a 7-day course of intravenous (IV) C/T (3 grams every 8 hours as 1-hour infusion) and inhaled polymyxin B. Two months prior to the current admission, the patient was admitted with recurrent pneumonia due to multidrug-resistant *P. aeruginosa* cultured from an endotracheal tube (ET) aspirate (saved as PA-NM-055, **Table 1**) that was treated with C/T and IV polymyxin B for 5 days. One month prior to the current admission, the patient was admitted with suspected pneumonia and urosepsis treated empirically with linezolid and aztreonam, with the latter agent changed to C/T plus inhaled polymyxin B for 5 days based on susceptibilities of the *P. aeruginosa* isolate cultured from the ET aspirate (isolate not saved - **Table 1**).

**Table 1.** Susceptibility profiles of *P. aeruginosa* isolates collected longitudinally from a single patient.

| Time of sample collection | Two months prior to admission |  | One month prior to admission | Hospital day 2 | Hospital day 16 |  |
| --- | --- | --- | --- | --- | --- | --- |
| Sample | ET<sup>b</sup> aspirate | ET<sup>b</sup> aspirate (repeat testing<sup>c</sup>) | ET<sup>b</sup> aspirate | Sputum | Bone biopsy<sup>d</sup> | Bone biopsy<sup>d</sup> (repeat testing<sup>c</sup>) |
| Isolate ID: | PA-NM-055 |  | Not available |  | PA-NM-086 |  |
| Antibiotic<sup>a</sup> / MIC (µg/mL) |  |  |  |  |  |  |
| Amikacin | ≥64 | 32 | ≥64 | ≥64 | 16 | 8 |
| Aztreonam | Resistant (KB) | 8 | Susceptible (KB) | Intermediate (KB) | Resistant (KB) | ≥32 |
| Cefepime | ≥64 | ≥32 | ≥64 | ≥64 | ≥64 | ≥32 |
| Ceftazidime | Resistant (KB) | ≥32 | Susceptible (KB) | Resistant (KB) | Resistant (KB) | ≥32 |
| Ciprofloxacin | ≥4 | ≥4 | ≥4 | ≥4 | ≥4 | ≥4 |
| Gentamicin | ≥16 | ≥16 | ≥16 | ≥16 | ≥16 | ≥16 |
| Meropenem | ≥16 | 8 | ≥16 | ≥16 | 8 | 4 |
| Piperacillin-tazobactam | Resistant (KB) | ≥128/4 | --<sup>e</sup> | Resistant (KB) | Resistant (KB) | ≥128/4 |
| Polymyxin B | 1 | 1<sup>f</sup> | --<sup>e</sup> | 0.75 | 1.5 | 1<sup>f</sup> |
| Tobramycin | ≥16 | ≥16 | ≥16 | ≥16 | ≥16 | ≥16 |
| Ceftolozane-tazobactam<sup>g</sup> | 0.75/4 | 0.75/4 | 0.75/4 | 0.5/4 | 192/4 | ≥256 |
| Ceftazidime-avibactam<sup>g</sup> | --<sup>e</sup> | 2/4<sup>c</sup> | 2/4 | 1.5/4 | 1.5/4 | 1/4<sup>c</sup> |

During the index admission, the patient experienced new fevers to 101°F (38.3°C), prompting an evaluation for infection, and was empirically treated with meropenem and linezolid. The patient was found to have methicillin-resistant *Staphylococcus aureus* (MRSA) growing from two of two blood cultures on hospital day 2 and sputum cultures grew *P. aeruginosa* (isolate not saved - **Table 1**). The patient was then switched from meropenem to C/T for possible pneumonia vs. tracheitis due to *P. aeruginosa,* and linezolid was changed to daptomycin for continuation of MRSA bacteremia treatment. After 7 days of *P. aeruginosa*-targeted treatment, the patient’s pulmonary symptoms improved, and C/T was discontinued; however, the patient continued to be intermittently febrile during the hospitalization. Diagnostic imaging studies suggested the cause of intermittent fevers was a new acute sacral osteomyelitis. On hospital day 16, a bone biopsy was performed, which grew ESBL-positive *Klebsiella pneumoniae*, *Morganella morganii*, MRSA, and *P. aeruginosa* (saved as PA-NM-086) demonstrating elevated minimum inhibitory concentrations (MICs) for meropenem (**Table 1**). The patient was continued on daptomycin, and meropenem (2 grams IV every 8 hours) was added as a continuous infusion for treatment of acute osteomyelitis. Additional susceptibility testing for C/T, CZA, and polymyxin B were requested. Four days after the biopsy, results of the MIC testing on PA-NM-086 revealed high-level C/T resistance but susceptibility to CZA (**Table 1**); therefore, IV meropenem was changed to IV CZA, IV polymyxin B, and metronidazole, with a planned 6-week course of therapy. On hospital day 85, the patient was discharged to an acute inpatient rehabilitation center after completing 6 weeks of IV antibiotics for osteomyelitis.

### Microbiology and Susceptibility Testing

Antibiotic susceptibility was tested for *P. aeruginosa* isolates collected from the patient across multiple admissions, comparator *P. aeruginosa* clinical isolates PS2045 and PS2046 collected previously from blood of patients hospitalized at NMH, and other *P. aeruginosa* strains constructed during the study. As part of routine patient care, antibiotic susceptibility testing for all clinical isolates from the patient was performed by the clinical microbiology laboratory using Vitek II (bioMérieux, Marcy l’Étoile, France), Kirby Bauer, or Etest (bioMérieux, Marcy l’Étoile, France or Liofilchem), depending on the drug. These results were verified for PA-NM-055 and PA-NM-086 by repeat testing in the lab (**Table 1**). All strains used in the study, including reference *P. aeruginosa* strains PA14 and PAO1, and *E. coli* strains used for cloning, are described in **Table S1**.

Confirmatory susceptibilities were determined for archived *P. aeruginosa* strains by microbroth dilution on Sensititre plates (Thermo Fisher Scientific). *P. aeruginosa* ATCC 27853 was used for quality control purposes. Briefly, isolates were grown overnight on Luria-Bertani (LB) agar at 37°C. The following day, 3-5 colonies were picked and re-suspended in water and diluted to an optical density at 600 nm (OD_600_) of 0.8-1.0. A total of 30 μL of the bacterial suspension was inoculated into 11 mL of cation-adjusted Mueller-Hinton Broth (CA-MHB). The positive control and wells containing antibiotic were inoculated with 50 μL of the suspension, and the negative control was inoculated with 50 μL of CA-MHB only. The plate was incubated at 37°C for 20 hours and read manually for growth. Antibiotic concentrations that led to >95% inhibition of growth (i.e. no visible growth) were noted as MICs.

C/T MICs for the clinical isolates and constructed strains were determined using antibiotic strips (bioMérieux, Marcy l’Étoile, France or Liofilchem). Bacterial strains were streaked onto LB agar plates and grown overnight at 37°C. A few individual colonies were resuspended in CA-MHB to achieve an OD_600_ of 0.2-0.3 that corresponded to 10^8^ CFU/mL. The bacterial suspension was plated onto cation-adjusted Mueller-Hinton agar (CA-MHA) using cotton swabs. A C/T antibiotic strip (Liofilchem) was placed on the plates and incubated for 16-20 hours at 37°C. Assessment of MIC was done as per the manufacturer’s instructions (i.e., the intersection of the strip resulting in >95% growth inhibition i.e. no visible growth was read as the C/T MIC).

### Whole Genome Sequencing

Whole genome sequencing (WGS) of the isolates PA-NM-055 and PA-NM-086 was performed as follows. Isolates from frozen stocks (−80°C) were streaked onto LB agar plates, and a single colony was cultured in LB broth overnight at 37°C. Genomic DNA was extracted using the Promega Maxwell 16 Cell DNA Purification Kit (Promega, Madison, WI, USA). Libraries were prepared using the Nextera XT DNA Library Preparation Kit (Illumina, Inc., San Diego, CA, USA). The clinical isolates were sequenced on a NextSeq 500 instrument (150 base pair reads, paired-end, mid-output flow cell, 133 M read pairs). Sequences were trimmed using Trimmomatic v0.32 (26) and then *de novo* assembly was performed with SPAdes 3.9.1 (27). Contigs shorter than 200 bp or with an average read coverage of less than 5x were excluded.

For long read sequencing of PA-NM-086, a sequencing library was prepared from unsheared and non-size-selected genomic DNA using the Oxford Nanopore ligation sequencing kit SQK-LSK110 and sequenced on a FLO-MIN106 flow cell on a MinION sequencing device. Guppy v3.4.5 was used to base call reads with the R9.4.1 high-accuracy model. Assembly of long reads was performed using the Trycycler v0.5.0 pipeline (28). Reads were filtered using Filtlong v0.2.1 to remove reads shorter than 1000 bases and the bottom 5% of the reads based on quality. Reads were then subsampled into 24 subsets using the Trycycler “subsample” function. Eight read subsets were assembled using Flye v2.8.3 (29), eight read subsets were assembled using Raven v1.5.3, and eight read subsets were assembled using minimap2 v2.21, miniasm v0.3, and minipolish v0.1.3 (30, 31). Clustering, reconciliation, circularization, multiple sequence alignment, and consensus sequence generation from the resulting 24 assemblies were performed using Trycycler. The consensus assembly was polished with medaka v1.4.3 using the “r941_min_hac_g507” model. Illumina reads were aligned to the assembly using BWA v0.7.17 (32), and assembly errors were corrected using Pilon v1.23 (33) with a minimum depth setting of 0.1.

Read alignment and Pilon correction were performed sequentially until no further assembly corrections were identified. Polypolish v0.4.3 (34) was used for final assembly correction using the Illumina short reads. Sequencing reads and genome assemblies are deposited in NCBI under BioProject accession PRJNA547625.

### Phylogenetic Analysis

To characterize the prevalence of L2 β lactamases among publicly available *P. aeruginosa* genome sequences, all *P. aeruginosa* genome assemblies (n=46,236, accessed Oct 14, 2025) in the NCBI assembly database were searched for the *bla*_L2_ gene. All sequence type 235 genomes (n=2996, accessed Oct 14, 2025) were downloaded and whole genome phylogenetic relationships were estimated using Mashtree v1.2.0 (35). The presence of the *bla*_L2_ gene sequence was identified in the genome sequences using nucleotide BLAST. Ten ST235 genome assemblies without a full-length *bla*_L2_ gene were selected randomly. Illumina read sequences were downloaded from NCBI for all L2-containing sequences identified by nucleotide BLAST against the NCBI WGS database with results limited to *P. aeruginosa* (taxid 287). For the four ST235 isolate sequences for which sequence reads were not available in the Sequence Read Archive (SRA) database of NCBI, pseudoreads were generated from the genome assemblies as previously described (36). Short read and pseudoread sequence alignment, phylogenetic analysis, and tree visualization were performed as previously described (37). Sequence reads and pseudoreads were aligned to the PA-NM-086 genome assembly.

### Construction of strains expressing bla_L2_

The *P. aeruginosa* reference strains PA14 and PAO1 were genetically modified to exogenously express *bla*_L2_ (**Table S1**). The gene *bla*_L2_ was amplified from the genomic DNA of PA-NM-086 using Q5 high fidelity polymerase (New England Biolabs) and primers 1 and 2 (**Table S2**). The amplicon was integrated at the EcoRI site downstream to the isopropyl-β-D-thiogalactopyranoside (IPTG)-inducible *lac*UV5 promoter in the plasmid pPSV37-Gen^R^ (38) by Gibson assembly (Thermo Fisher Scientific). The resulting recombinant plasmid (designated “p*bla*_L2_”) was introduced into the conjugative *Escherichia coli* strain SM10 (39) by calcium chloride heat shock method and the transformants were selected on LB agar containing gentamicin (15 µg/mL). Conjugation was performed to transfer the recombinant plasmid from the transformant SM10 strain to the *P. aeruginosa* strains, and the exconjugants were selected on LB agar containing gentamicin (30 µg/mL) and irgasan (triclosan, 5 µg/mL). The plasmid construct was isolated (Qiagen) from the exconjugant *P. aeruginosa* strains and verified by Sanger sequencing using internal primers recognizing the *bla_L2_* gene and an adjacent plasmid pPSV37 sequence (**Table S3**). The vector control strains were constructed by introducing the empty vector pPSV37 into PA14 and PAO1 using the same method. These strains were grown on CA-MHA containing gentamicin (15 µg/mL) with or without 1 mM IPTG for determining C/T MICs.

### RNA extraction and reverse-transcriptase-quantitative-PCR (RTqPCR)

Overnight cultures of bacterial strains grown in LB medium were subcultured in LB medium and grown to an OD_600_ of 0.4. IPTG (1 mM) and imipenem (1 µg/mL) were added to test for the induction of gene expression in the indicated experiments. Negative control cultures were left untreated. All bacterial cultures were then incubated for an additional 2 hours. A volume of bacterial culture corresponding to 1 x 10^8^ CFU was centrifuged, and the pellet was resuspended in RNA protect reagent (Qiagen). After incubation for 5 min at room temperature, samples were centrifuged again, and bacterial cells in the pellet were treated with lysozyme (15 µg/mL in Tris-EDTA). RNA was extracted using an RNEasy kit (Qiagen), and cDNA was synthesized with Superscript IV VILO reverse transcriptase kit (Invitrogen) using master mix with and without reverse transcriptase, the latter used as a negative control to assess for DNA contamination. RT-qPCR was performed by using qPCR probe assays (Integrated DNA Technologies) on a CFX-96 real-time thermocycler (Biorad). Expression level was determined by normalizing the threshold cycle (Ct or Cq) value of each target gene (Ct_target_) for every sample to that of the respective housekeeping gene *rpoD* (Ct_reference_) generating ΔCt values and plotted as fold change in gene expression with respect to the mean ΔCt of multiple biological replicates of the control sample. Each experiment was carried out at least twice (i.e., at least 2 biological replicates). The primers and probes used for the RT-qPCR are listed in **Table S2**.

### Generation of deletion mutants

Gene deletion mutants in the comparator *P. aeruginosa* strain PS2045 background were generated using a suicide-vector mediated allelic replacement method (40). Fwd-up and Rev-up primers were used to generate a ∼500 bp fragment homologous to 5’ end (‘up’ fragment), whereas Fwd-down and Rev-down primers were used to generate a ∼500 bp fragment homologous to 3’ end (‘down’ fragment) of the target gene (**Table S3**). However, a 936 bp transposon gene immediately 3’ to the *bla*_L2_ gene in PS2045 is duplicated elsewhere in the genome of PS2045, preventing amplification of an ‘up’ fragment. To avoid the duplicated region, we targeted deletion of only a 477 bp internal segment of *bla*_L2_ instead of the entire gene. This segment encodes the S-X-X-K motif and SDN loop, which are required for the activity of the L2 β-lactamase (41). The ‘up’ and ‘down’ fragments were combined using the Fwd-up and Rev-down primers to generate the full fragment, which was integrated at the HindIII site of the plasmid pex18-Hyg^R^ (40) by Gibson assembly (Thermo Fisher). The resulting plasmid was introduced into the conjugative *E. coli* strain SM10 (39) by calcium chloride heat shock method. The transformants were selected on LB agar containing hygromycin (100 µg/mL) and used for conjugation with the strain PS2045 at 37°C for 1 hour. The resulting PS2045 merodiploids with the entire plasmid integrated into the genome were selected using LB agar supplemented with hygromycin (500 µg/mL) and irgasan (5 µg/mL). These merodiploids were resolved by counter-selection on LB agar supplemented with sucrose (10 ug/mL) to select for mutants that lost the *sacB-*containing vector backbone along with the target gene. The deletion of the target gene in the selected colonies was verified by PCR amplification with confirmatory primers (**Table S3**) followed by Sanger sequencing of the PCR product. Genomic DNA from mutant isolates was sequenced using a commercial service (SEQCENTER) to produce paired-end 150 bp reads. Sequencing reads were aligned to the complete genome assembly of isolate PS2045 and assessed for mutations relative to the reference using Breseq v0.39.0 (42).

### Statistical analysis

Each experiment was carried out at least twice and results were plotted using GraphPad prism version 10. Statistical significance of the difference observed between datasets, i.e. the median MICs or the mean fold changes in gene expression, was determined by one-way ANOVA with Tukey’s multiple comparisons test for comparing multiple samples and Student’s t test for comparing 2 samples, generating *p* values, where *p* < 0.05 was considered statistically significant.

## Results

### Antibiotic resistance phenotypes of P. aeruginosa isolates collected from the patient

The respiratory isolate collected 2 months prior to the index admission (PA-NM-055) and the respiratory isolate collected on day 2 of the index admission were both susceptible to C/T and CZA (**Table 1**). In contrast, the bone biopsy isolate collected on hospital day 16 (PA-NM-086) was found to be highly resistant to C/T while remaining susceptible to CZA, giving the clinical impression that C/T resistance emerged after treatment with repeated courses of C/T. However, genomic analysis revealed that PA-NM-055 belonged to the sequence type (ST) 111, whereas PA-NM-086 belonged to ST235, indicating that the patient had experienced infections by at least two distinct high-risk clones of *P. aeruginosa* over time. Due to the limited number of isolates available for analysis, we were unable to determine whether PA-NM-086 represented a new infection that contributed to acute osteomyelitis observed in the patient or whether the PA-NM-086 lineage had been present in this patient for an extended period and at some point replaced the more susceptible PA-NM-055 after multiple treatment courses.

### Characterization of antimicrobial resistance (AMR) genes in the C/T resistant strain PA-NM-086

To investigate possible mechanisms of C/T resistance in PA-NM-086, we performed WGS. Long read *de novo* assembly with short read correction yielded a single circular 6.92 Mb chromosome. Interestingly, mutations known to be associated with C/T resistance in *P. aeruginosa* (15–24) were absent in PA-NM-086. In particular, mutations in the class C β-lactamase gene *bla*_AmpC_, which encodes the Pseudomonas-derived cephalosporinase (PDC) member of the AmpC family, were not observed in PA-NM-086. The AmpC family of beta-lactamases are involved in peptidoglycan recycling and are commonly associated with C/T resistance (4, 16, 21–24, 43–53). However, an Ambler Class A β-lactamase gene *bla*_L2_ with 100% identity with the L2 β-lactamase gene found in many *Stenotrophomonas maltophilia* strains was identified (41). This gene was not present in the respiratory isolate PA-NM-055, which was C/T sensitive.

### Frequency of bla_L2_ in P. aeruginosa

We next examined whether *bla*_L2_ was present in other *P. aeruginosa* strains. We first searched a publicly available *P. aeruginosa* whole-genome sequence repository (∼46,000 genomes) for other ST235 isolates that contained the *bla*_L2_ gene. Only six of the 2,995 ST235 genomes examined contained this gene, suggesting that the presence of the *bla*_L2_ gene in *P. aeruginosa* is quite rare even within this high-risk clonal lineage (**Fig. 1A**). Across all the *P. aeruginosa* genomes (∼46,000) available in NCBI, *bla*_L2_ was identified in 12 isolates, 11 of which were collected in the USA (**Fig 1B**). Further, this gene was predominantly confined to just 5 lineages (**Fig. 1B**), suggesting that the transfer of the *bla*_L2_ gene from *S. maltophilia* to *P. aeruginosa* was a globally rare event.

**Fig. 1.**
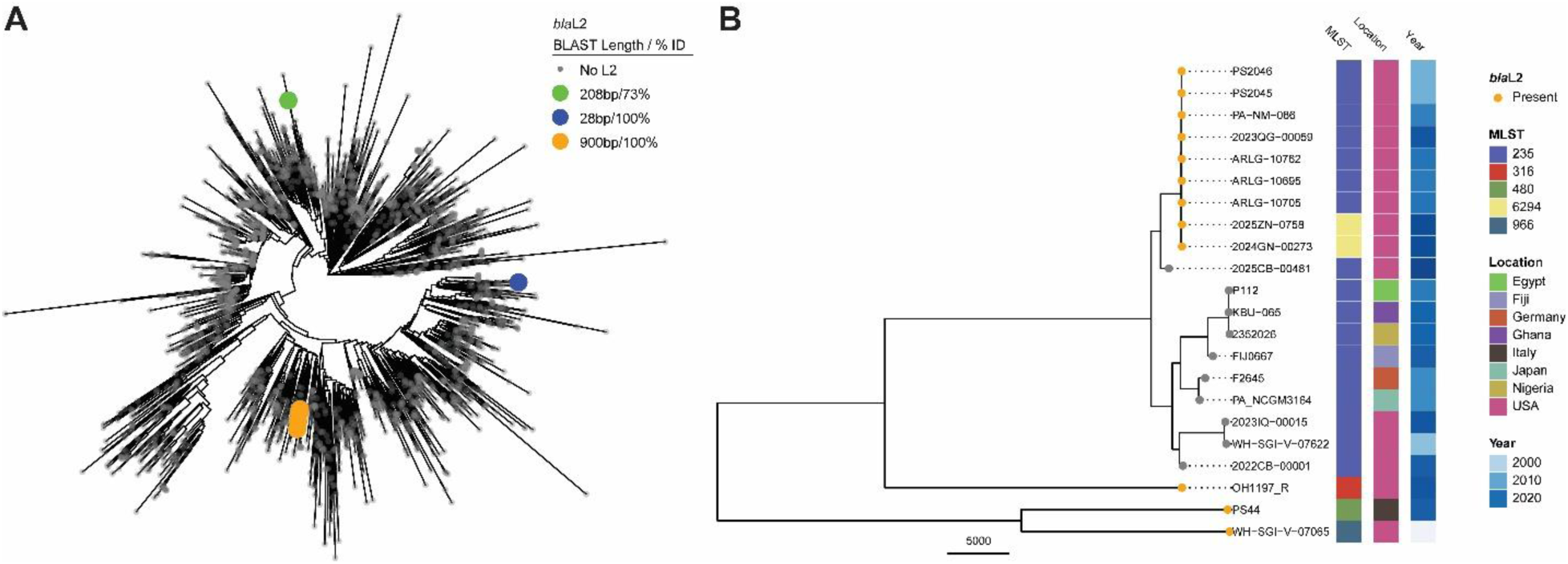
The presence of the *bla* gene among publicly available *P. aeruginosa* genomes. **A)** Whole-genome phylogenetic analysis of ST235 genome assemblies in NCBI (n=2995, one outlier removed) using Mashtree. Isolates with blastn hits for the *bla*_L2_ gene are highlighted. Taxa are colored by the alignment length and percent identity of the BLAST results. Six genomes had identical genes (orange), and two genomes contained portions of the *bla*_L2_ gene (blue and green). **B)** Phylogenetic relationships of all *bla*_L2_-positive genome sequences in NCBI by Mashtree analysis. Isolates carrying *bla*_L2_ gene (orange) with > 88% blastn sequence identity across > 95% length of the *bla*_L2_ gene and relative to the 10 randomly selected non-*bla*_L2_ ST235 isolates from NCBI (gray) are shown. Bars show the multilocus sequence type (MLST), geographic location and decade of collection of each isolate.

### Genetic context of bla_L2_

We used the complete genome sequence of PA-NM-086 to determine the genomic context of *bla*_L2_. In *S. maltophilia,* the *bla*_L2_ gene is adjacent to the *ampR*_L2_ gene, which encodes a transcriptional regulator of *bla*_L2_ (54). In PA-NM-086, we found that the 912 bp *bla*_L2_ gene is adjacent to an identical *ampR*_L2_ gene except that the length of the *ampR*_L2_ gene is reduced from 867 bp to 507 bp (**Fig. 2**). We designated this truncated *ampR*_L2_ allele as “*ampR_L2_*^tr^” that encodes an open reading frame of 168 amino acids of which the first 129 amino acids are identical to that of AmpR_L2_ of *S. maltophilia*. Of note, the genomes of nearly all *P. aeruginosa* isolates contain an *ampR* gene (here designated as “*ampR*_AmpC_”) directly adjacent to *bla*_AmpC_ (53), which is distinct from *ampR*_L2_ in *S. maltophilia* (71.6% identity over 67% of the *ampR*_AmpC_ gene). The truncated *ampR*_L2_^tr^ gene in PA-NM-086 has only 72% identity over 41% of *ampR*_AmpC_. These findings indicate that *ampR*_L2_^tr^ likely originated from *S. maltophilia* rather than from gene duplication of *ampR*_AmpC_. The genes *bla*_L2_ and *ampR*^tr^ are part of a larger 9,034 bp element that contains 7 other genes and is repeated in tandem 5 times in the genome of PA-NM-086 (**Fig. 2**). Additional AMR genes identified in this element include two tetracycline resistance genes [*tetR* and *tetA,* coding for the tetracycline repressor protein class G and tetracycline efflux pump, respectively (55, 56)] and an aminoglycoside resistance gene [*aph*, coding for an aminoglycoside 3’ phosphotransferase (57)]. This genomic region contains IS91 family transposases, suggesting that it is part of a composite transposon or insertion sequence (IS)-associated resistance cassette.

**Fig. 2.**
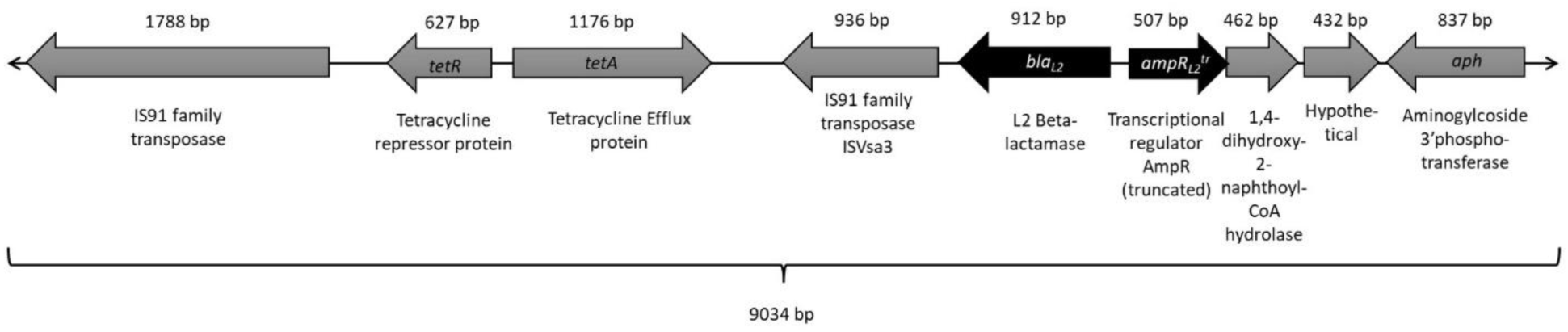
Genomic element containing *bla*_L2_. Five tandem copies of the genomic element (9,034 bp) shown in the figure were found in PA-NM-086. The length and annotation of each open reading frame is indicated.

### Impact of exogenous expression of the L2 β-lactamase on C/T resistance in P. aeruginosa

Since *S. maltophilia* strains are relatively resistant to C/T (58–60) and *bla*_L2_ was originally found in *S. maltophilia* (61, 62), we hypothesized that the gain of *bla*_L2_ contributed to C/T resistance in the strain PA-NM-086. To test this hypothesis, we constructed a plasmid p*bla*_L2_ to exogenously express *bla*_L2_ gene in the presence of IPTG and introduced it in the *P. aeruginosa* reference strains PA14 and PAO1. We compared the C/T MICs in the presence and absence of IPTG and found that the addition of IPTG increased the C/T MIC in PA14-p*bla*_L2_ (**Fig. 3**) and PAO1-p*bla*_L2_ (**Fig. S1**) by approximately four-fold (i.e., two doubling dilutions), which was statistically significant. In contrast, the addition of IPTG had no effect on C/T resistance in strains containing the empty vector (PA14-EV and PAO1-EV).

**Fig. 3.**
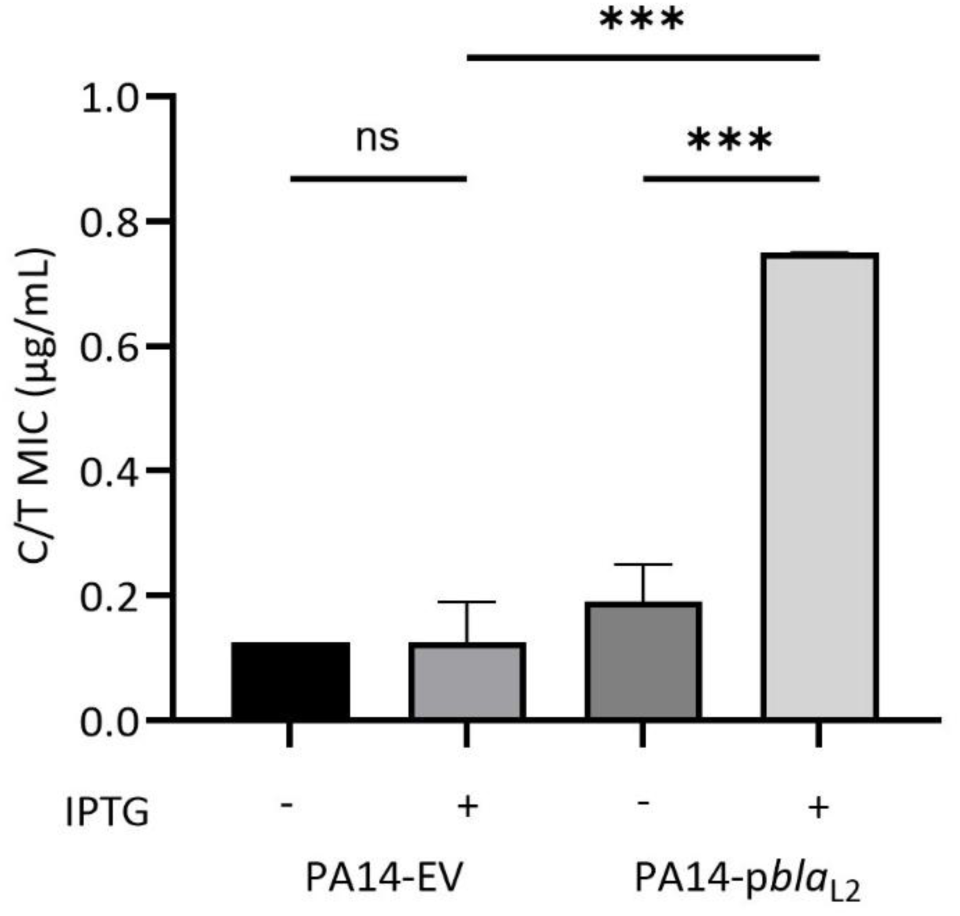
Effect of exogenous expression of *bla*_L2_ in *P. aeruginosa* strain PA14 on C/T MIC. C/T MICs are shown for the strains PA14-EV and PA14-*pbla*_L2_ with (+) or without (−) IPTG as determined by antibiotic strip tests. The bars represent median MIC values of 3 biological replicates with confidence intervals shown as error bars. One-way ANOVA was used to determine statistical significance, indicated as *p* values, where ns = *p* >0.05 and *** = *p* <0.001. Abbreviations-EV, empty vector; MIC, minimal inhibitory concentration; IPTG, isopropyl-β-D-thiogalactopyranoside.

### Associations between the bla_L2_ expression and C/T resistance in P. aeruginosa

We next examined whether differences in *bla_L2_* gene copy number were associated with different degrees of C/T resistance. As mentioned, PA-NM-086 contains five copies of the *bla_L2_* gene. A search of *P. aeruginosa* WGSs in our collections identified 2 clinical isolates, PS2045 and PS2046 (**Table S1**), containing 1 and 2 copies, respectively, of the 9,034 bp *bla*_L2_-containing genetic element (**Fig. 1B**). We determined the C/T MICs of these two isolates. Whereas PS2045 had a low C/T MIC of ∼4 µg/mL, PS2046 was highly resistant to C/T, with an MIC of >256 µg/mL, consistent with a relationship between *bla*_L2_ gene copy number and MIC.

We reasoned that strains with higher *bla*_L2_ copy numbers may express more L2 β-lactamase, which would in turn lead to higher C/T MICs. To examine this, we performed RT-qPCR on PA-NM-086, PS2045, and PS2046. The expression levels of the *bla*_L2_ genes in these strains correlated with *bla*_L2_ copy number (**Fig. S2A**). Notably, expression of *bla*_L2_ in PA14-p*bla*_L2_ under uninduced (C/T MIC ∼0.19 µg/mL) and induced (C/T MIC ∼0.75 µg/mL) conditions correlated with C/T resistance of PA14-p*bla*_L2_ in these conditions (**Fig. 3** and **Fig. S2A**). However, expression of *bla*_L2_ in the clinical isolates PS2045 (C/T MIC ∼4 µg/mL) and PS2046 (C/T MIC >256 µg/mL) was lower than that in strain PA14-p*bla*_L2_ in the presence of IPTG induction (C/T MIC ∼0.75 µg/mL). This suggested that the expression level of *bla*_L2_ was not solely responsible for C/T MICs in these strains. Because over-expression of *bla*_AmpC_ has been linked to C/T resistance (16), we measured *bla*_AmpC_ expression but found similar *bla*_AmpC_ transcript levels in all strains (**Fig. S2B**). We also found that the sequences of *bla*_AmpC_ gene in PS2045, PS2046 and PA-NM-086 are identical. Further, known mutations associated with increased C/T resistance in *P. aeruginosa* were absent in PS2045 and PS2046 like PA-NM-086. Therefore, difference in C/T resistance levels among these strains do not appear to be related to the inherent levels of *bla*_AmpC_ expression in these strains.

Overall, these findings indicate that exogenous expression of a *bla*_L2_ gene leads to a modest but statistically significant increase in resistance to C/T in *P. aeruginosa* and that other mechanisms may augment this resistance.

### Loss of L2 β-lactamase decreases C/T resistance in P. aeruginosa

To confirm the role of L2 β-lactamase in C/T resistance in PA-NM-086, we sought to examine the effect of deletion of the *bla_L2_* gene on C/T MICs. This was difficult in PA-NM-086 because it contains five copies of the *bla*_L2_ gene. Since PS2045 contains only one copy of *bla*_L2_ (**Fig. 2**), it allowed the targeted deletion of the *bla*_L2_ gene. We deleted approximately 50% of the *bla*_L2,_ open reading frame, including the region coding for the active site of L2 β-lactamase, to construct the mutant strain PS2045Δ*bla_L2_*^act^ (see Methods). The C/T MIC value of PS2045Δ*bla*^act^ (∼0.75 µg/mL) was significantly lower than that of the parental strain (∼4 µg/mL) (**Fig. 4**). This indicated that a functional L2 β-lactamase contributes to a modest level of C/T resistance in *P. aeruginosa*.

**Fig. 4.**
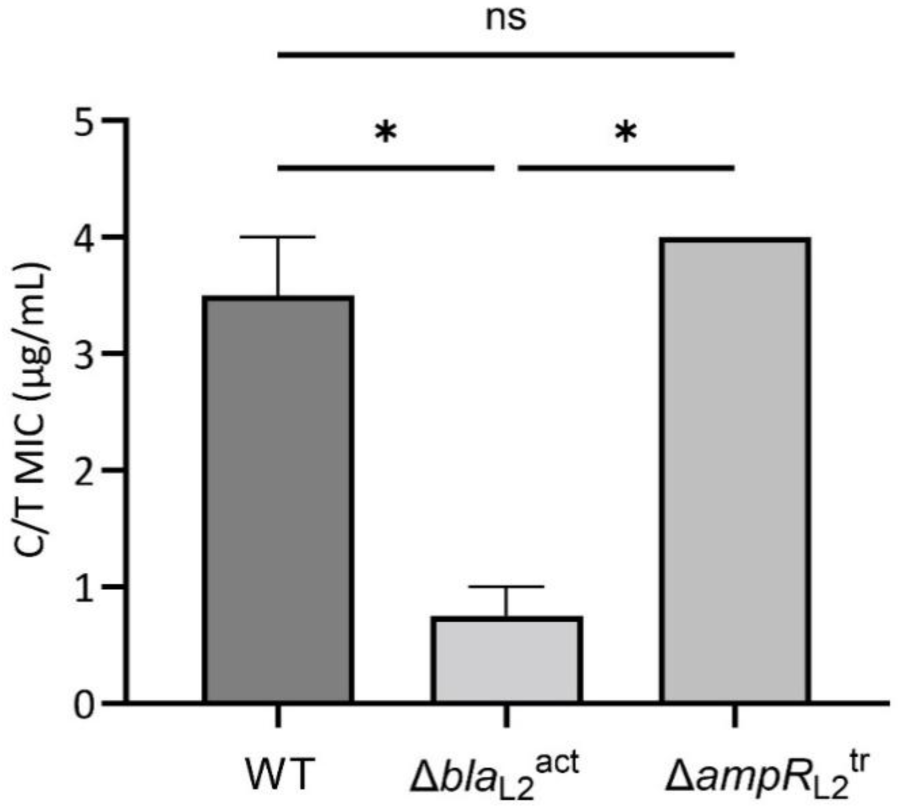
C/T MICs for a parental PS2045 strain compared to PS2045 strains with disruptions of the *bla*_L2_ or *ampR*_L2_^tr^ genes. C/T MICs in the parental (WT) and mutant strains of the clinical isolate PS2045 were determined by antibiotic strip tests. Disruption of the *bla*_L2_ gene (Δ*bla*_L2_^act^) significantly reduced C/T MICs, whereas deletion of the *ampR*_L2_^tr^ gene (Δ*ampR*_L2_^tr^) had no effect. The bars represent median MIC values from 2 biological replicates, and error bars represent confidence intervals. One-way ANOVA was used to determine statistical significance, indicated as *p* values, where ns = *p* >0.05 and * = *p* <0.05.

### L2 β-lactamase in P. aeruginosa has activity against other β-lactam antibiotics

The L2 β-lactamase of *S. maltophilia* has broad activity against β-lactam antibiotics (41, 61, 63, 64). We assessed the effect of disruption of *bla*_L2_ on the resistance of the strain PS2045 to other β-lactams. The mutant strain PS2045Δ*bla*^act^ displayed reduced levels of resistance to aztreonam, cefepime, ceftazidime and piperacillin/tazobactam in comparison to the parental strain (**Table 2**). However, this was not observed for the carbapenems imipenem, meropenem, and doripenem, as the parental PS2045 strain was susceptible to all three carbapenems despite the presence of an intact *bla*_L2_ gene. The antibiotic resistance profiles of the clinical isolates examined in this study with one, two, and five copies of the *bla*_L2_ gene (PS2045, PS2046, PA-NM-086, respectively) were also consistent with a role for the L2 β-lactamase in resistance to multiple β-lactams (**Table 2**). These findings demonstrated that the L2 β-lactamase increases detectable resistance to multiple β-lactams but probably not to carbapenems.

**Table 2.**
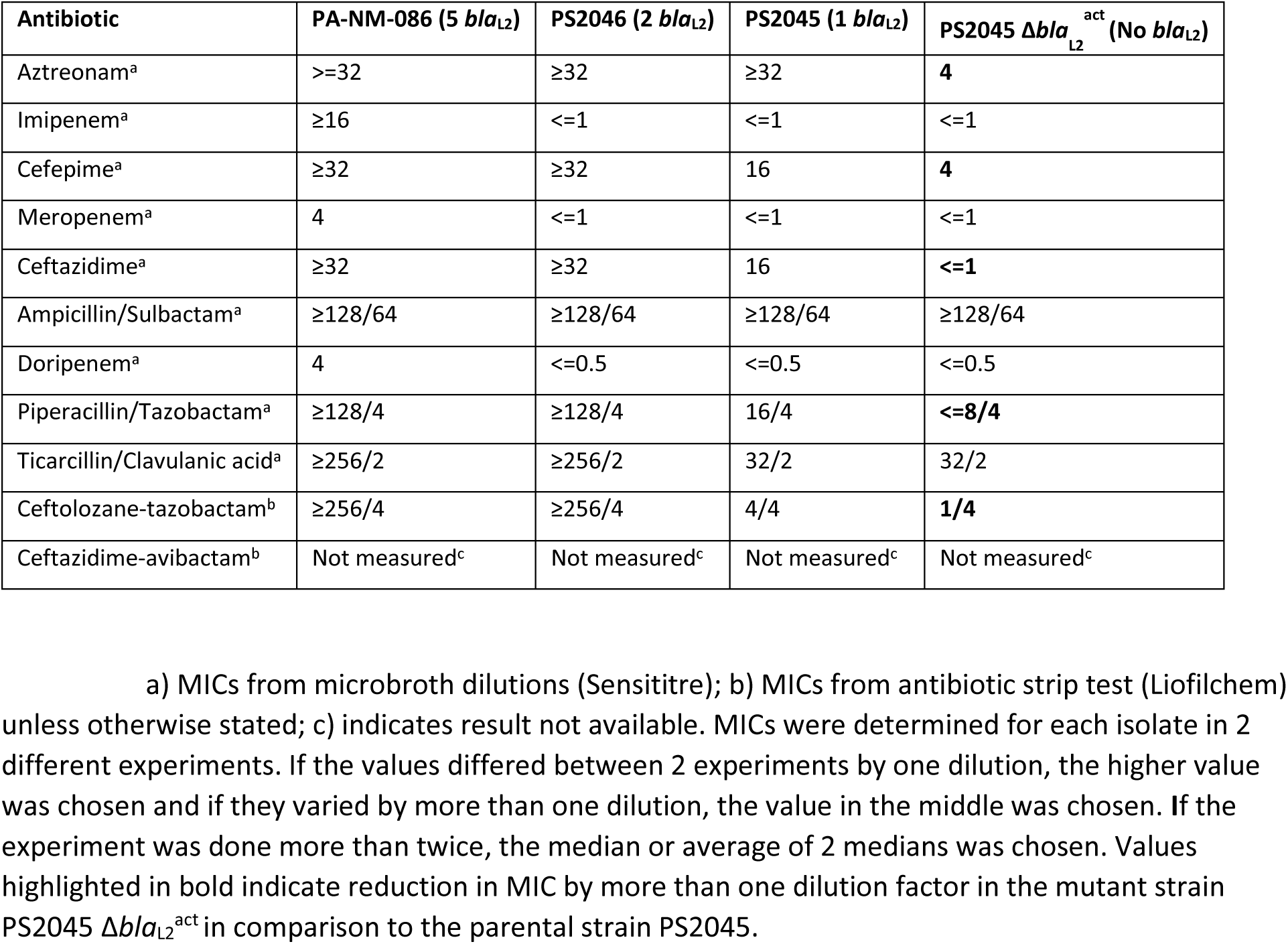
Susceptibility profiles of *P. aeruginosa* isolates containing different copy numbers of *bla*_L2_ and the mutant containing a disrupted *bla*_L2_ gene (PS2045Δ*bla*_L2_^act^).

### AmpR_L2_^tr^ does not regulate expression of the L2 β-lactamase in PA-NM-086

In *S. maltophilia,* AmpR_L2_ regulates expression of *bla*_L2_, and induces *bla*_L2_ expression in the presence of the β-lactam antibiotic imipenem (54). As *ampR_L2_*^tr^ is substantially truncated (867 bp reduced to 507 bp) in PA-NM-086, we hypothesized that *bla*_L2_ expression in this strain may not be regulated by AmpR^tr^. To test this, we determined the expression levels of *bla*_L2_ in PA-NM-086 in the presence and absence of a low concentration of imipenem (1 µg/mL). We used PA14-p*bla*_L2_ strain as a negative control because *bla*_L2_ expression in this strain is controlled by IPTG induction of the *lac*UV5 promoter (**Fig. S2A**). As a positive control, we examined the expression of *bla*_AmpC_, which has been reported to be induced by imipenem (65). As expected, *bla*_L2_ expression in PA14-p*bla*_L2_ was induced by IPTG and not imipenem (**Figs. S2A and 5A**), and *bla*_AmpC_ was induced by imipenem in PA-NM-086 as well as PA14-p*bla*_L2_ (**Fig. 5B**). In PA-NM-086, no significant difference was observed in *bla*_L2_ expression levels in the presence or absence of imipenem (**Fig. 5A**). This indicated that *bla*_L2_ expression in PA-NM-086 occurs constitutively, consistent with the interpretation that *ampR_L2_*^tr^ does not modulate *bla_L2_* expression in the absence of imipenem. We also generated a deletion mutant of *ampR*^tr^ in PS2045 and found that deletion of *ampR*^tr^ did not affect the C/T MIC of PS2045 (**Fig. 4**). This agreed with the lack of AmpR_L2_^tr^-dependent regulation of *bla_L2_* expression in PA-NM-086 (**Fig. 5A**). Taken together, these results suggest that L2 β-lactamase contributes to C/T resistance in *P. aeruginosa* independently of the product of the *ampR_L2_*^tr^ gene.

**Fig. 5.**
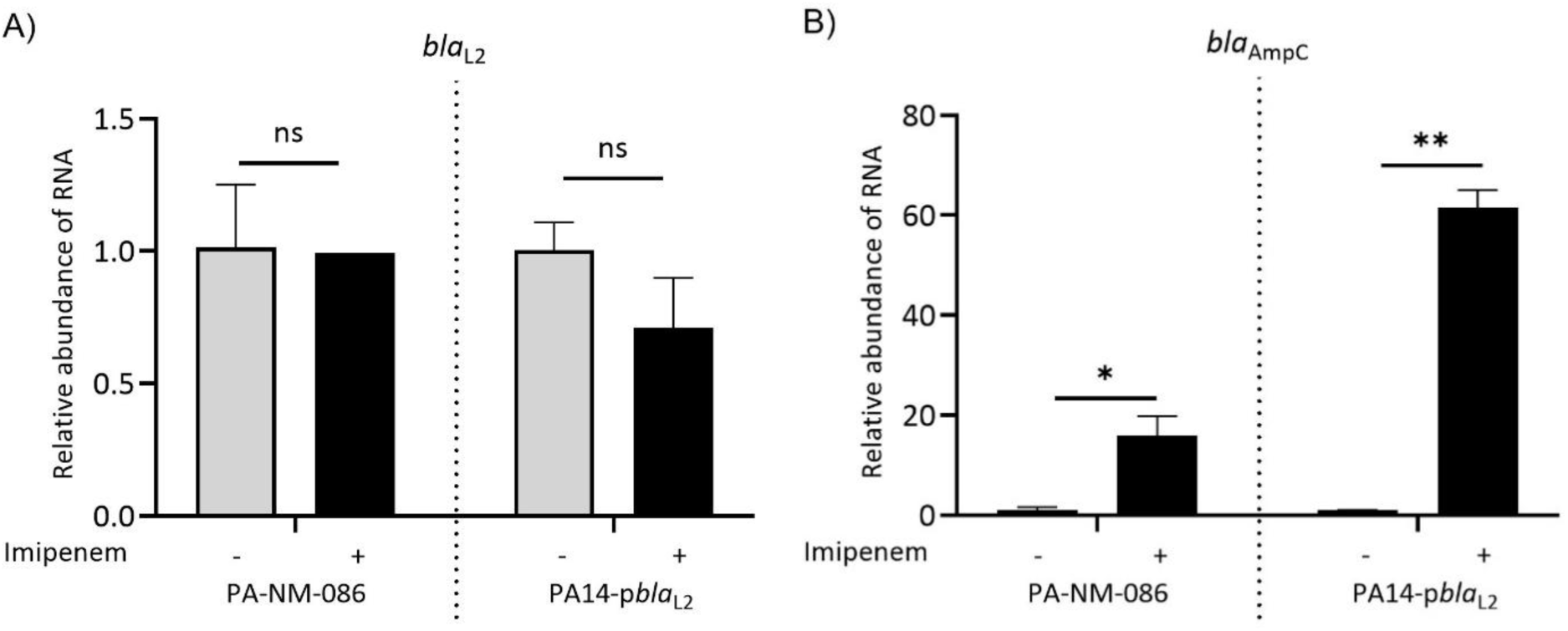
Expression of *bla*_L2_ and *bla*_AmpC_ in the presence and absence of imipenem. Expression of the genes **A)** *bla*_L2_ and **B)** *bla*_AmpC_ in the strains PA-NM-086 and PA14-*pbla*_L2_ in the presence (+) or absence (−) of imipenem was determined by RT-qPCR. Gene expression is shown as relative abundance in RNA levels in comparison to the respective untreated control (−) after normalizing to the house keeping gene *rpoD*. The bars represent means of 2 biological replicates. Error bars represent standard deviations. Student’s t test was used to determine statistical significance, indicated as *p* values, where ns = *p* >0.05, * = *p* <0.05, and ** = *p* <0.01.

## Discussion

Our investigation suggests that the L2 β-lactamase gene, usually associated with *S. maltophilia*, has been transferred to *P. aeruginosa* on more than one occasion, though this remains a globally rare event. The L2 β-lactamase, along with L1 β-lactamase, is produced by many *S. maltophilia* strains and is largely responsible for the widespread resistance of this species to most β-lactams, including third-generation cephalosporins (61, 66, 67). *S. maltophilia* demonstrates C/T MIC range of 16/4 to >128/4 µg/mL (58–60). The L2 β-lactamase may contribute to this relatively high level of C/T resistance. In our study, we found that introduction of *bla*_L2_ into the *P. aeruginosa* strains PA14 and PAO1 caused a modest approximately four-fold increase in C/T MICs. Likewise, disruption of the L2 β-lactamase encoding gene in PS2045 resulted in an approximate four-fold reduction in the C/T MIC. The L2 β-lactamase is therefore yet another example of cross-species transfer of an antibiotic resistance mechanism. Although PS2045, PS2046, and PA-NM-86 are very closely related, they have 1, 2, and 5 copies of the ∼9 kb element, respectively, suggesting that selective pressures exist that cause repeated duplication of this element.

The bone-biopsy isolate (PA-NM-086) had 5 copies of the *bla*_L2_ gene and a C/T MIC >256 µg/mL, which matched with C/T MIC of a blood isolate, PS2046, that contained only 2 copies of *bla*_L2_. In contrast, the blood isolate, PS2045, containing only one copy of *bla*_L2_, displayed a low MIC of ∼4 µg/mL, which suggests non-linear scaling of resistance (as measured by MIC) as copy number of *bla*_L2_ gene increases or that the C/T MICs of these strains are influenced by additional resistance determinants. We hypothesize that exposure to C/T concentrations in excess of 4X MIC may result in eradication of strains with 1 copy of *bla*_L2_. Such PK/PD exposures are achievable for strains with C/T MICs of 1 µg/mL (68), but would be much more difficult to achieve for strains with an MIC of 4 µg/mL or greater and even more so when site-of-infection exposures are lower than those attained in plasma with maximal labeled dosing (69).

The 100 percent identity of the *bla*_L2_ gene sequences in *P. aeruginosa* and *S. maltophilia* suggests that the former acquired the gene from the latter. The gene appears to have been obtained from *S. maltophilia* along with a truncated *ampR*_L2_ gene as part of a 9,034 bp genetic element. L2, a class A serine β-lactamase, is encoded by a chromosomal gene in *S. maltophilia* (41). Chromosomally encoded genes are less readily transferred across species than those encoded on plasmids, which may explain why *bla*_L2_ was identified in only a handful of *P. aeruginosa* genomes. Both *P. aeruginosa* and *S. maltophilia* are ubiquitous environmental organisms, found in soil, plant material, rivers, lakes, and streams (70, 71), which would provide ample opportunities for genetic exchange. Interestingly, both bacteria are also commonly cultured from the lungs of individuals with cystic fibrosis (72), which may be another site at which genetic exchange between these bacteria occurred.

This is the second report of the presence of an L2 β-lactamase gene in *P. aeruginosa* but appears to represent a more recent transfer of the gene. A previous report by Tian and colleagues described in *P. aeruginosa* the acquisition of the *bla*_PME-1_ gene, which is similar to *bla*_L2_ (73). However, several observations suggest that the *bla*_PME-1_ gene is distinct from *bla*_L2_. While the L2 β-lactamase of PA-NM-086 has 100% amino acid identity to the corresponding protein of *S. maltophilia,* PME-1 has only 50% amino acid identity (73). (This is substantially less than the 68% amino acid identity observed between SHV-1 and TEM-1, which are 2 distinct β-lactamases.). The *bla*_PME-1_ gene is carried on a 9-kb plasmid and was not adjacent to an *ampR*_L2_ gene. Although PME-1 conferred resistance to third-generation cephalosporins including ceftazidime, it was inhibited by tazobactam and would therefore be predicted to be inactive against C/T. We therefore suggest that PME-1 and L2 β-lactamase be considered distinct β-lactamases.

Several lines of evidence indicate that the L2 β-lactamase contributes to C/T resistance. First, multiple *bla*_L2_-containing isolates of *P. aeruginosa* had elevated MICs to C/T. Second, upon exogenous expression of *bla*_L2_ in two distinct *P. aeruginosa* strains (PA14 and PAO1) that did not naturally contain it, we observed a modest but significant increase in the C/T MIC levels. Third, the disruption of *bla*_L2_ in PS2045 caused a modest but significant reduction in C/T MICs. Together, these findings support the contribution of *bla*_L2_ to C/T resistance in *P. aeruginosa*. A number of exogenously acquired β-lactamases can enhance the resistance of *P. aeruginosa* to C/T, including GES-6, SHV-2a, VIM-2, and IMP-28 (49, 51, 74, 75). Our findings indicate that the L2 β-lactamase should be added to this list.

Our findings suggest that the L2 β-lactamase contributes to C/T resistance independently of the *ampR*^tr^ gene in *P. aeruginosa*. In *S. maltophilia,* AmpR_L2_-dependent induction of *bla*_L2_ expression occurs in the presence of β-lactam antibiotics such as imipenem (54). However, in the absence of β-lactam antibiotics, some expression of *bla*_L2_ can occur independently of AmpR_L2_ in *S. maltophilia* (76). Therefore, it is possible that the *P. aeruginosa* strains used in this study, which naturally contain *bla*_L2_ gene, express *bla*_L2_ constitutively irrespective of the presence of an inducer antibiotic and independently of regulation by AmpR_L2_^tr^. In *S. maltophilia,* induction of *bla_L1_* and *bla_L2_* is dependent on AmpR_L2_; disruption of the regulator results in reduced activity of the L2 β-lactamase and corresponding reductions in the MICs of several β-lactams (54). We observed that in the presence of AmpR_L2_^tr^, L2 expression was not affected by the inducer imipenem, suggesting a loss of the context-dependent repressor and activator effects typically associated with AmpR (77). Although our study shows that L2 β-lactamase contributes to C/T resistance independently of a functional AmpR_L2_ in the strain PS2045, it remains to be tested whether the presence of a functional AmpR_L2_ would further influence L2 β-lactamase-dependent C/T resistance in *P. aeruginosa*. AmpR_L2_ has been suggested as a potential drug target due to its role in resistance to multiple β-lactams (78); however, since L2 β-lactamase contributes to C/T resistance even in the absence of regulation by AmpR_L2_, such an approach would be ineffective against L2-expressing strains of *P. aeruginosa*.

To date, much work on C/T resistance in *P. aeruginosa* has been focused on the core genome chromosomal *bla*_AmpC_ gene, a class C β-lactamase (16, 17, 21, 24, 44–47, 50). Mutations leading to *bla*_AmpC_ de-repression can increase the MIC of C/T to 16 ug/mL for *P. aeruginosa* (75). The range of mutations that predispose clinical strains to overproduce *bla*_AmpC_ remains incompletely understood (79). We found that *bla*_AmpC_ in PA-NM-086 was induced in the presence of the β-lactam imipenem, suggesting that it could also be contributing to C/T resistance. A second mechanism of *bla*_AmpC_ mediated resistance to C/T is the structural alteration of AmpC itself. Paired isolate studies (comparing isolates collected pre- and post-C/T treatment) have found that alterations within the *bla*_AmpC_ open reading frame are major drivers of increased C/T MICs and phenotypic resistance (17, 80). *P. aeruginosa* isolates with a deletion of a 21 bp region of the *bla*_AmpC_ have increased C/T MIC values of >256 µg/mL (4). Additionally, alterations within the omega-loop of AmpC appear to widen the enzyme’s binding pocket, leading to increased ceftolozane and ceftazidime hydrolysis and reduced affinity for avibactam (50). However, the *bla*_AmpC_ gene present in PA-NM-086 did not exhibit any of the mutations associated with C/T resistance, making it unlikely that structural changes in AmpC were directly responsible for the elevated C/T MIC in this strain.

Our study has several limitations. First, we were not able to ascertain what portion of PA-NM-086 C/T resistance was due to the L2 β-lactamase. Our MIC analysis indicated that the five copies of the *bla*_L2_ gene in PA-NM-086 may cause higher C/T MICs, as strains with more copies of the *bla*_L2_ gene had higher C/T MICs. Because we were unable to delete all five copies of this gene in PA-NM-086, we could not verify this. However, the non-linear impact of *bla*_L2_ copy number on C/T resistance level observed in the *P. aeruginosa* strains used in this study (zero to one: 2 doubling dilutions and one to two copies: > 2 doubling dilutions) suggests that there is an association between *bla*_L2_ gene copy numbers and C/T MICs in our isolates. Confirmatory studies are needed. Second, we lacked an antecedent C/T-susceptible strain of the same lineage as PA-NM-086 from our patient to strengthen a comparative genomics analysis. Third, our copy number experiments and deletion-expression analyses used different strains of *P. aeruginosa,* so we were unable to take into account the effects of other strain background differences on C/T resistance. Fourth, the induction of expression of *bla*_L2_ in PA14 and PAO1 did not result in the same C/T MIC as observed in PS2046, which expressed *bla*_L2_ lower than PA14-p*bla*_L2_ with IPTG induction. However, the log-scale changes in C/T MIC were similar upon exogenous expression and deletion of *bla*_L2_, suggesting that the L2 β-lactamase can have a measurable impact on C/T resistance even at low copy numbers.

## Conclusion

In conclusion, mutational and expression analyses suggest that acquisition of the class A β-lactamase gene *bla*_L2_ contributed to C/T resistance in *P. aeruginosa*. Additional studies are needed to better understand the processes that select for the transfer of resistance genes between *P. aeruginosa* and *S. maltophilia*. This study reports a rare but important case of antibiotic resistance that extends our understanding of C/T resistance in *P. aeruginosa*.

## Acknowledgements

This work was supported in part through the computational resources and staff contributions provided by the Genomics Compute Cluster, which is jointly supported by the Feinberg School of Medicine, the Center for Genetic Medicine, and Feinberg’s Department of Biochemistry and Molecular Genetics, the Office of the Provost, the Office for Research, and Northwestern Information Technology. The Genomics Compute Cluster is part of Quest, Northwestern University’s high-performance computing facility, with the purpose to advance research in genomics. The core facility for Sanger Sequencing at Northwestern University and ACGT DNA sequencing services are acknowledged for carrying out Sanger sequencing of PCR products.

## Funding

Research reported in this publication was supported by the National Institute of Allergy and Infectious Diseases/National Institutes of Health grants R01AI118257 (ARH) and R01AI158530 (NJR). The content is solely the responsibility of the authors and does not necessarily represent the official views of the National Institutes of Health.

## Potential conflicts of interest

Dr. O’Donnell received funding from Merck and Co., Inc. that owns C/T. All other authors declare no conflict of interest.

## Author Contributions

*Concept and Design: Garai, Hauser, Rhodes*

*Acquisition, analysis, or interpretation of data: Garai, Ozer, Nozick, Jozefczyk, Hauser, Nam, O’Donnell, Rhodes*

*Drafting of the initial manuscript: Garai, Jozefczyk, Rhodes*

*Critical revision of the manuscript for important intellectual content: Garai, Ozer, Nozick, Jozefczyk, Hauser, Nam, O’Donnell, Rhodes*

*Data Analysis: Rhodes, Ozer, Garai Obtained Funding: Rhodes, Hauser*

*Administrative, technical, or material support: Nozick, O’Donnell*

*Supervision: Rhodes, Ozer, Hauser*

## Data availability

The source data associated with the manuscript are available on the Prism repository of Northwestern University and can be accessed through the digital object identifier (doi) https://doi.org/10.18131/8g2gz-7zt62.

## Notes

https://prism.northwestern.edu/records/8g2gz-7zt62

### Summary of Updates

Added funding source, external data doi, and supplementary data.

