## Supplementary material for "A novel mechanism of ceftolozane-tazobactam resistance in *Pseudomonas aeruginosa* mediated by L2 β-lactamase": Fig S1 and S2; Tables S1 S2 and S3

**Affiliations:** ^1^Department of Microbiology-Immunology, Northwestern University, Feinberg School of Medicine, Chicago, IL, USA; ^2^ Midwestern University College of Pharmacy Downers Grove Campus, Downers Grove, IL and Department of Pharmacy, Northwestern Medicine, Chicago, IL, USA; ^3^Stanford University School of Medicine, Stanford CA, USA; ^4^School of Medicine, University of California Irvine, Irvine, CA, USA; ^5^Department of Pharmacy Practice, Albany College of Pharmacy and Health Sciences, Albany, NY, USA; ^6^Division of Infectious Diseases, Department of Medicine, Northwestern University Feinberg School of Medicine, Chicago, IL, USA; ^7^Department of Pharmacy Practice, Midwestern University College of Pharmacy Downers Grove Campus, Downers Grove, IL, USA; ^8^Midwestern University College of Pharmacy Downers Grove Campus, Pharmacometrics Center of Excellence, Downers Grove, IL, USA; ^9^Department of Pharmacy, Northwestern Medicine, Chicago, IL, USA.

**#Corresponding author and reprints:** Nathaniel J. Rhodes, PharmD, MSC, BCPS AQ-Infectious Diseases, Associate Professor of Pharmacy Practice. Department of Pharmacy Practice, Midwestern University College of Pharmacy Downers Grove Campus. 555 31st Street, Downers Grove, IL. 60515; Office: AH364, 630-515-7376; Fax: 630-515-6958;

***Present address:** Department of Pharmacy, Prisma Health – Greenville Memorial Hospital, Greenville, South Carolina, USA

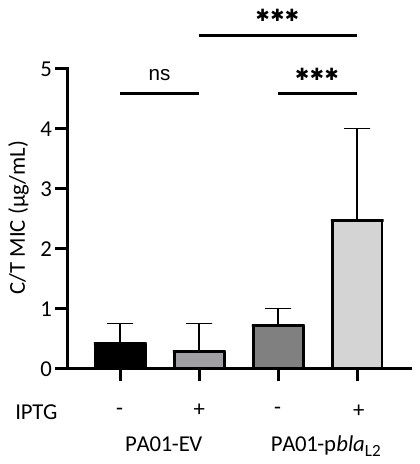

**Fig. S1. Effect of exogenous expression of *bla*_L2_ in *P. aeruginosa* strain PAO1 on C/T MICs.** C/T MICs for the strains PAO1-EV and PAO1-*pbla_L2_* with (+) or without (-) IPTG determined by antibiotic strip tests. The bars represent median MIC values of 4 biological replicates with confidence intervals shown as error bars. One-way ANOVA was used to determine statistical significance, indicated as *p* values, where ns = *p* >0.05 and *** = *p* <0.001. Abbreviations- EV, empty vector; MIC, minimal inhibitory concentration; IPTG, isopropyl-β-D-thiogalactopyranoside.

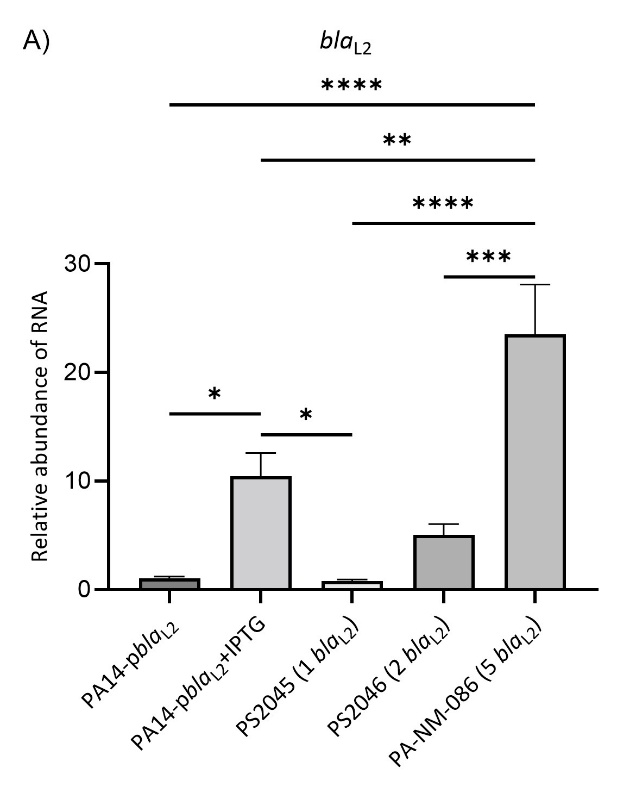

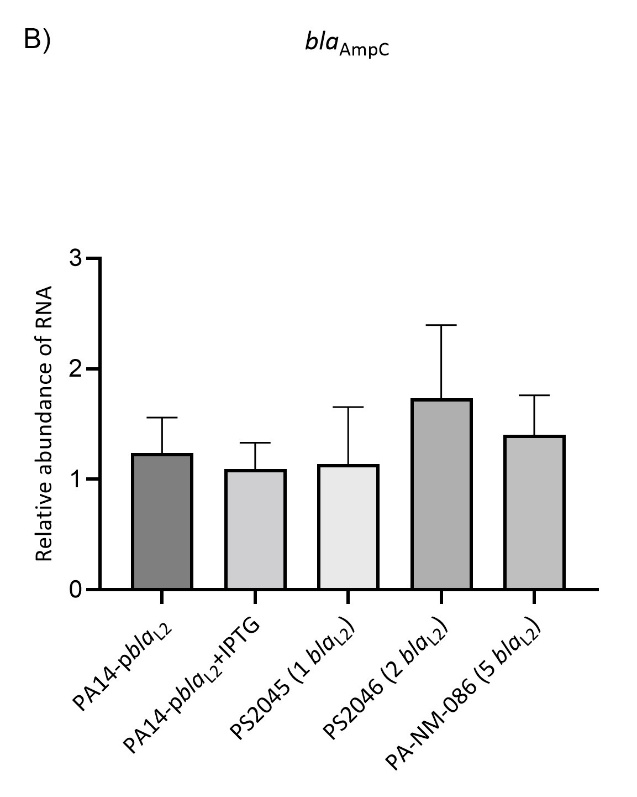

**Fig. S2. *bla*_L2_ expression in strains containing different *bla*_L2_ copy numbers.** Expression of the genes **A)** *bla*_L2_ and **B)** *bla*_AmpC_ in the strains PA14-p*bla*_L2_ (with and without IPTG), PS2045, PS2046 and PA-NM-086 was determined by RT-qPCR. The number of *bla_L2_* genes is indicated in parentheses. Gene expression is shown as relative abundance in RNA levels with respect to PA14-p*bla*_L2_ without IPTG after normalizing to *rpoD*. The bars represent mean of at least 2 biological replicates. Error bars represent standard deviation. One-way ANOVA was used to determine statistical significance and indicated as *p* values, * = *p* <0.05, ** = *p* <0.01, *** = *p* <0.001, **** = *p* <0.0001. Abbreviations- IPTG, isopropyl-β-D-thiogalactopyranoside.

**Table S1. Bacterial strains used in this study.**

| **Species** | **Strain** | **Characteristics** | **Source (Reference)** | **GenBank Accession number** |
| --- | --- | --- | --- | --- |
| *P. aeruginosa* | PA-NM-055 | Collected from endotracheal tube aspirate | This study, NMH | JBUYFB000000000.1 |
| *P. aeruginosa* | PA-NM-086 | Collected from bone biopsy | This study, NMH | JBUYFC000000000.1 |
| *P. aeruginosa* | PS2045 | Collected from blood | This study, NMH | JAHTGK000000000.2 |
| *P. aeruginosa* | PS2046 | Collected from blood | This study, NMH | JAHTGL000000000.2 |
| *P. aeruginosa* | PA14 | Reference strain |  | GCA_000014625.1 |
| *P. aeruginosa* | PA14-pPSV37 (empty vector) | Lab construct, PA14 containing pPSV37 plasmid | This study | - |
| *P. aeruginosa* | PA14-p*bla*_L2_ | Lab construct, PA14 containing p*bla*_L2_ plasmid | This study | - |
| *P. aeruginosa* | PAO1 | Reference strain |  | GCA_000006765.1 |
| *P. aeruginosa* | PAO1-pPSV37 (empty vector) | Lab construct, PAO1 containing pPSV37 plasmid | This study | - |
| *P. aeruginosa* | PAO1-p*bla*_L2_ | Lab construct, PAO1 containing p*bla*_L2_ plasmid | This study | - |
| *P. aeruginosa* | PS2045 ∆*bla*_L2_^act^ | Lab construct | This study | SRR37233608 |
| *P. aeruginosa* | PS2045 ∆*ampR*_L2_^tr^ | Lab construct | This study | SRR37233606 |
| *E. coli* | Top10 | F- *mcrA* Δ(*mrr*-*hsdRMS-mcrBC*) φ80*lacZ*ΔM15 Δ*lacX74 nupG recA1*  *araD139* Δ(*ara-leu*)7697 *galE15 galK16 rpsL*(StrR) *endA1* λ- | Invitrogen | - |
| *E. coli* | SM10  λpir | TpR SmR *recA*, *thi*, *pro*, *hsdR*-M+RP4: 2-Tc:Mu: Km Tn7 λpir | (1) | - |

**Footnotes:** Bacterial strains used in the study are listed along with their characteristics and sources. GenBank accession numbers for the strains used in this study are provided. References are provided in parentheses. Abbreviations- NMH, Northwestern Memorial Hospital, Chicago.

**Table S2.** **RTqPCR primers and probes.**

| **Primer/Probe** | **Sequence** |
| --- | --- |
| *rpoD*_qPCR FWD | 5’GGGCGAAGAAGGAAATGGT 3’ |
| *rpoD*_qPCR REV | 5’CTGGATCAGGTCGAGGAATTG 3’ |
| *rpoD*_qPCR PRB | 5’TCCATCGCCAAGAAGTACACCAACC 3’ |
| *bla*_L2__qPCR FWD | 5’ATCATCACCAGCGACAACAC 3’ |
| *bla*_L2__qPCR REV | 5’CCCTTGGCAAAGCTGTTCA 3’ |
| *bla*_L2__qPCR PRB | 5’AACCTGCTGTTCGGCGTGGT 3’ |
| *bla*_AmpC_ qPCR FWD | 5’GATGCTCGGGTTGGAATAGAG 3’ |
| *bla*_AmpC_ qPCR REV | 5’TCGACCTCGCGACCTATAC 3’ |
| *bla*_AmpC_ qPCR PRB | 5’TGCAGAAGGACCAGGCACAGATC 3’ |

**Footnotes:** Forward (FWD) and Reverse (REV) primers, and probes (PRB) used for determining expression of the genes *rpoD*, *bla*_L2_ and *bla*_AmpC_ are listed with their sequences.

**Table S3. Primers and plasmids used for cloning and generating gene deletions are listed.**

| **Primer** | **Sequence** |
| --- | --- |
| pPSV37 EcoRI *bla*_L2_ Fwd | 5’ gattacgaattgtagctagctaggATGCTCGCCCGTCGCCGATTCC 3’ |
| pPSV37 EcoRI *bla*_L2_ Rev | 5’ gaggatccccgggtaccgagctcgTCATCCGATCAACCGGTCG 3’ |
| *bla*_L2_ internal Fwd | 5’ GCGACAACACCGCCGCCAACC 3’ |
| *bla*_L2_ internal Rev | 5’ GCGGGCGGGCCACCGACCACG 3’ |
| pPSV37 internal Fwd | 5’ CAGGAAACAGCTATGACCAT 3’ |
| pPSV37 internal Rev | 5’ GCGATCAAAAAACCCCTCAA 3’ |
| PS2045 *bla*_L2_^act^ Fwd up | 5’ gtaaaacgacggccagtgccaGGATGTATGCGCGCGCCGAACC 3’ |
| PS2045 *bla*_L2_^act^ Rev up | 5’ aactcgagccgcaagcatgctgaaGCAGTGACCGCGTTCCTGC 3’ |
| PS2045 *bla*_L2_^act^ Fwd down | 5’ ttcagcatgcttgcggctcgagttCTTGATCGGGGGCGACCAGC 3’ |
| PS2045 *bla*_L2_^act^ Rev down | 5’ gagtcgacctgcaggcatgcaCTCATCGCGCGACCCTAGC 3’ |
| PS2045 *bla*_L2_^act^ KO confirmatory Fwd | 5’ GTTGGCCAGTTCCGTCAGTTGC 3’ |
| PS2045 *bla*_L2_^act^ KO confirmatory Rev | 5’ GCGCACATAGTTGGCGTGATAG 3’ |
| PS2045 *ampR*_L2_^tr^ Fwd up | 5’ gtaaaacgacggccagtgccaGTGACCGGTGCATGCGAGAGC 3’ |
| PS2045 *ampR*_L2_^tr^ Rev up | 5’ aactcgagccgcaagcatgctgaaGCTGGAGTGATGGATGATCC 3’ |
| PS2045 *ampR*_L2_^tr^ Fwd_down | 5’ ttcagcatgcttgcggctcgagttCGACCACCTGTTGCCGGTGC 3’ |
| PS2045 *ampR*_L2_^tr^ Rev down | 5’ gagtcgacctgcaggcatgcaGCAGCAGGATCGTGACGATC 3’ |
| PS2045 *ampR*_L2_^tr^ KO confirmatory Fwd | 5’ GGTGTTGTCGCTGGTGATGATC 3’ |
| PS2045 *ampR*_L2_^tr^ KO confirmatory Rev | 5’ CTTGGCCGCATCGGTCCAGC 3’ |
| **Plasmid** | **Characteristics (Reference)** |
| pPSV37-Gen^R^ | Exogenous expression vector, Gen^R^, *colE1* origin, PA origin, *oriT*, *lacUV5* promoter, *lacIq*, stops in every reading frame preceding the MCS and T7 terminator following the MCS (relative to the lacUV5 promoter) (2) |
| pex18-Hyg^R^ | Allelic exchange vector, Hyg^R^, *oriT*, *sacB*, *lacZα*, MCS from pUC18 (3) |

**Footnotes**: **Primers** used for cloning indicate the plasmid and restriction site in their names and those used for gene deletion indicate the strain name and the target sequence. Cloning was verified with internal primers and gene deletion was verified with knockout (KO) confirmatory primers (see methods). All primer sequences have the region homologous to the gene amplified highlighted in capital letters. **Plasmid** names are shown with their marker antibiotic resistance genes (Gen- Gentamicin; Hyg- Hygromycin) and characteristics of the plasmids are shown with the source (reference) of the plasmid.
